# A Bayesian approach to incorporate structural data into the mapping of genotype to antigenic phenotype of influenza A(H3N2) viruses

**DOI:** 10.1101/2022.03.26.485931

**Authors:** William T. Harvey, Vinny Davies, Rodney S. Daniels, Lynne Whittaker, Victoria Gregory, Alan J. Hay, Dirk Husmeier, John W. McCauley, Richard Reeve

**Author notes:** RIP 28^th^ September 2019.

## Abstract

Surface antigens of pathogens are commonly targeted by vaccine-elicited antibodies but antigenic variability, notably in RNA viruses such as influenza, HIV and SARS-CoV-2, pose challenges for control by vaccination. For example, influenza A(H3N2) entered the human population in 1968 causing a pandemic and has since been monitored, along with other seasonal influenza viruses, for the emergence of antigenic drift variants through intensive global surveillance and laboratory characterisation. Statistical models of the relationship between genetic differences among viruses and their antigenic similarity provide useful information to inform vaccine development though accurate identification of causative mutations is complicated by highly correlated genetic signals that arise due to the evolutionary process. Here, using a sparse hierarchical Bayesian analogue of an experimentally validated model for integrating genetic and antigenic data, we identify the genetic changes in influenza A(H3N2) virus that underpin antigenic drift. We show that incorporating protein structural data into variable selection helps resolve ambiguities arising due to correlated signals, with the proportion of variables representing haemagglutinin positions decisively included, or excluded, increased from 59.8% to 72.4%. The accuracy of variable selection judged by proximity to experimentally determined antigenic sites was improved simultaneously. Structure-guided variable selection thus improves confidence in the identification of genetic explanations of antigenic variation and we also show that prioritising the identification of causative mutations is not detrimental to the predictive capability of the analysis. Indeed, incorporating structural information into variable selection resulted in a model that could more accurately predict antigenic assay titres for phenotypically-uncharactrised virus from genetic sequence. Combined, these analyses have the potential to inform choices of reference viruses, the targeting of laboratory assays, and predictions of the evolutionary success of different genotypes, and can therefore be used to inform vaccine selection processes.

## Introduction

Antigenic variation is a mechanism by which an infectious agent such as a virus or bacterium alters the proteins or carbohydrates exposed to the host immune system to allow escape from immunity conferred by prior infection with or vaccination against a related agent. Antigenic drift seen in influenza viruses is a prime example of this process. Human seasonal influenza epidemics are estimated to infect around 15% of the global population annually resulting in three to five million cases of severe illness and in the order of 290,000 to 650,000 deaths annually (1,2). Influenza viruses evolve rapidly, with the frequency of mutations being highest in the genes encoding the surface glycoproteins such that antigenic variants can emerge. Some antigenic variants may rise to dominance among circulating viruses due to strong immune-mediated positive selection favouring viruses that infect individuals previously immune due to prior infection or vaccination. Consequently, human seasonal influenza vaccines must be frequently updated to ensure the antigenic responses they elicit will be active against viruses in circulation.

Seasonal influenza epidemics are caused by viruses belonging to two influenza A subtypes, A(H1N1) and A(H3N2), and by influenza B viruses which are classified into antigenically distinct lineages, B/Victoria and B/Yamagata. Trivalent vaccines contain antigens based on the influenza A(H1N1) and A(H3N2) and the predominant influenza B lineage while quadrivalent compositions include antigens representative of both influenza B lineages. To monitor the genetic and antigenic evolution of human influenza viruses, the WHO coordinates the Global Influenza Surveillance and Response System (GISRS) collaborating with academic scientists and national public health organisations (3,4). Vaccine strain selection is based on the antigenic and genetic evolution of circulating influenza viruses throughout the year and recommendations are developed by representatives of the GISRS at twice-yearly vaccine composition meetings.

Within each of the influenza A subtypes and influenza B lineages that cause seasonal influenza epidemics, the global virus populations typically consist of several antigenically distinct groups of viruses. It takes around six months to develop, produce and deliver an updated influenza vaccine so strain selection decisions must be made up to nine months in advance of the period when influenza viruses will circulate in a forthcoming season. It is therefore necessary to understand the antigenic similarity of circulating viruses to current vaccine strains and vaccine candidates to predict which antigenic variants are most likely to circulate at high frequency in advance of a future influenza season, a task that benefits from predictive modelling (5). Various approaches have demonstrated prediction of successful influenza lineages, from those emerging, with potential to inform vaccine virus selections (6–8). One approach is to use the shape and branching pattern of haemagglutinin (HA) phylogenetic trees to track and extrapolate changes in genotype frequency (6). Another approach is to predict lineage fitness using counts of amino acid substitutions inside and outside described antigenic sites as proxies for antigenic drift and reduced stability respectively (7); an approach that can be adapted to incorporate data from assays used to measure antigenic drift. Both the accuracy of and the reliance on such predictive models depend on an understanding of the indirect link between genotype and reproductive fitness in a partially immune population, a relationship that is informed by the more direct phenotypic consequences of genetic changes.

The antigenic characterisation of circulating viruses is dependent upon haemagglutination inhibition (HI) and virus neutralisation (VN) assays, both of which are used to assess the antigenic similarity of a circulating test virus to a panel of reference viruses that includes previous and current vaccine viruses and other candidate vaccine viruses. The panel of reference viruses, and post-infection ferret antisera raised against them, are selected to represent the diversity of antigenic phenotypes observed over the most recent seasons. Various modelling approaches have used data from antigenic assays to quantify similarity (9), and to explore the relationship between genetic and antigenic evolution allowing predictions of antigenic relationships from sequence data (8,10–12). A general challenge for modelling genotype-phenotype relationships is differentiating causative mutations from those that are non-causative and correlate with phenotypic changes due to genetic hitchhiking. One technique to improve differentiation is to explicitly account for the shared evolutionary history of viruses. Various comparative methods already exist to account for shared evolutionary history, but these tend to focus on traits intrinsically associated with particular taxa rather than ones that relate to relationships between taxa, as is the case here when working with pairwise measures of antigenic similarity. However, by including terms that represent branches of the phylogenetic tree, it is possible to account for shared evolutionary history and to prevent false statistical support for genetic terms due to repeated measurements (13).

Comparing sequences of test and reference viruses, while accounting for phylogenetic correlations, we have previously identified substitutions responsible for antigenic evolution among human influenza A(H1N1) and avian influenza A(H9N2) viruses (12,14). We then made this phylogenetically-aware model more statistically rigorous within a Bayesian framework with antigenic determinants identified using ‘spike-and-slab’ priors (15), a method of variable selection (SABRE) demonstrated to outperform alternative approaches such as LASSO (16) and elastic net regularised regression (17) as well as our own previous work (12). We further extended these approaches (eSABRE) by the inclusion of variables representing the underlying HI titre for each reference/test virus pair (18). However, in order to allow us to fully characterise the algorithms involved, the data analysed in that study comprised only 43 viruses. These previous approaches can therefore be characterised as either applying methodologically-limited methods of model selection to datasets of scientific relevance (12), or the application of state-of-the-art Bayesian approaches to datasets of limited scale and relevance to the problem of antigenically characterising viruses (18).

Here we develop these techniques into a practical tool for antigenic characterisation of viruses using Bayesian stochastic search variable selection (BSSVS) within a hierarchical model structure applied to a large set of antigenic data spanning 25 years and consisting of over 38,000 individual antiserum titres derived using panels of antisera and contemporary circulating test viruses. The BSSVS approach for model selection, which allows us to identify the combination of genetic variables best explaining the observed antigenic variation, is performed within the model as it undergoes Markov chain Monte Carlo (MCMC) sampling. Variables under investigation are free to drop out and return to the model as the optimal combination of terms is converged upon. Consequently, parameter estimates are made averaging over the best set of models, allowing uncertainty in the optimal combination of variables to be accounted for directly. A further advantage of a Bayesian approach here is that existing knowledge can be used to define the prior distribution of parameters. These priors, together with the likelihood of the observed data given the statistical model, combine using standard Bayesian methods to form the posterior distribution of parameters. We show that information derived from measurements of solved protein structures can be used to shape prior distributions and improve the accuracy with which we can attribute changes in antigenic phenotype to causative amino acid substitutions. Finally, using Bayesian model averaging, where predictions are averaged over a range of the best supported models (19), these approaches show the ability to accurately predict antigenic relationships from genetic sequences, whilst actively aiming to optimise variable selection.

## Modelling approach

The approach presented infers the genetic determinants of antigenic evolution by attributing variation in antigenic assays to differences in the amino acid sequences of reference and test viruses, while accounting for both phylogenetic structure in the data and other non-antigenic factors that cause variation in titres. Using fixed quantities of reference and test viruses, commonly eight or four haemagglutinating units in a particular HI assay, titres are recorded as the reciprocal of the maximum dilution of antiserum raised to a particular reference strain that is able to inhibit agglutination of red blood cells by a test virus. Lower titres, expressed as fold-drops, therefore reflect reduced antigenic similarity. The log_2_ titre is modelled reflecting the two-fold serial dilution of antiserum in assays. We describe below how the variation in titres attributable to the antigenic properties of viruses can be attributed to virus HA-gene sequences, firstly by mapping antigenic changes to branches of the phylogenetic tree and secondly by attributing antigenic changes to specific amino acid differences between reference and test viruses. Model selection was performed using BSSVS via binary indicator variables associated with each branch or amino acid position (18). These indicator variables, also known as binary mask variables, take the value zero or one determining variable exclusion (masking) or inclusion, respectively. The optimal combination of branches, or amino acid positions at which substitutions explain antigenic changes, is therefore determined by sampling these binary mask variables using MCMC.

Throughout this approach, we model the assay titre *Y* measured for an antiserum raised against reference virus *r* and each specific virus *v* on a given date *d* as lognormally distributed:

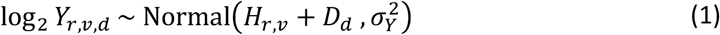

The log_2_ titre has a mean determined by combining the underlying log_2_ titre for each combination of reference virus and test virus, *H_r,v_*, and an effect accounting for day-to-day experimental variability in titres, *D_d_*. The use of log_2_ reflects the use of a two-fold serial dilution of antiserum, with recorded titres being the reciprocals of these dilutions. Residual variance of measured titres around this mean is represented by 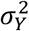. To improve the succinctness and therefore clarity of this section, prior distributions and other implementation details are provided in the *Model Implementation* subsection of the *Materials and Methods*, and details of indices, parameters, variances and other minor terms are described in Table 1.

**Table 1.**
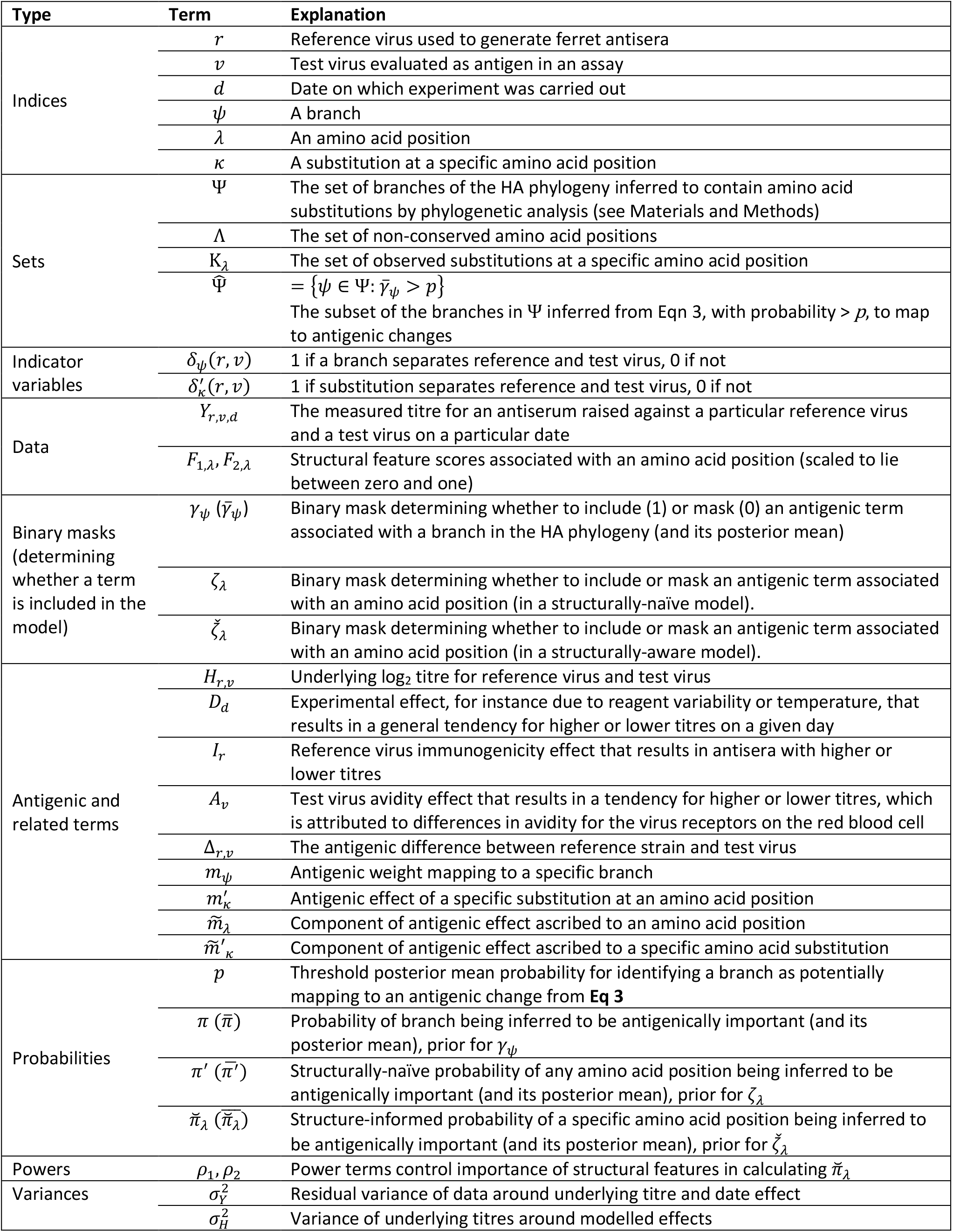
Model indices, terms and parameters.

The underlying log_2_ titre in **Eq 1** is itself normally distributed:

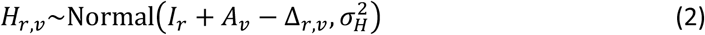

Underlying titres are modelled as depending on three general characteristics of the assayed viruses and antisera. The contributions of effects for reference strain immunogenicity, *I_r_*, test virus avidity, *A_v_*, and the antigenic relationship, Δ*_r,v_* are each inferred. The immunogenicity and avidity terms reflect general reactivity of antisera and viruses respectively. These manifest as a trend for higher or lower titres against all viruses or antisera for which titres are measured, independent of antigenic relationships. The genetic determinants of this antigenic component is of principal interest, so in the remainder of this section we describe how variation in this term is attributed to differences in the HA protein (see Table 1 for further details on the other effects). Since antigenic differences between reference and test viruses manifest as lower titres, the antigenic component of the model is constrained to take only non-negative values and is subtracted from the other terms in the model. Δ*_r,v_* is defined in several different ways in Eqs 3, 4, 5, 6 and 9, below, to allow us to compare different models of this antigenic relationship.

Antigenic difference is initially modelled as a linear combination of effects that occur during the phylogenetic evolution of the assayed viruses, with terms representing every branch, *ψ*, of the phylogeny to which amino acid substitutions were mapped in an ancestral state reconstruction (see Materials and Methods), Ψ, tested as predictors of reduced HI titres:

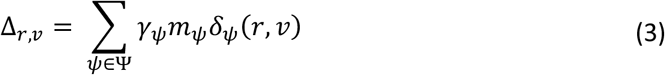

For each branch, a precomputed indicator variable *δ_ψ_*(*r, v*) is one when the branch falls on a direct path through the tree separating the reference and test viruses, and zero otherwise. Consequently, the titre for each combination of reference virus and test virus only depends on the antigenic weights for the combination of branches that fall between them on a path traced through the tree. The parameter *m_ψ_* is the antigenic weight associated with the branch, the expected drop in log_2_ HI titre when two viruses are separated in the phylogeny by branch *ψ*. The binary mask variable, *γ_ψ_*, takes the value zero or one determining whether branch *ψ* is either excluded from or included in the model, respectively, and each antigenic effect, *m_ψ_*, represents the antigenic effect of a specific branch when it is included (*γ_ψ_* = 1). When *γ_ψ_* is zero, any antigenic weight attributed to the branch is nullified (as the product, *γ_ψ_m_ψ_*, is zero). A higher proportion of MCMC samples with *γ_ψ_* = 1 indicates higher support in the data for an antigenic change mapping to branch *ψ*. For each branch, the proportion of *γ_ψ_* = 1, which is also the posterior mean value, 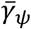,is referred to as the inclusion probability for the branch. This allows antigenic changes in the evolution of assayed viruses to be mapped to specific branches of the phylogeny to generate 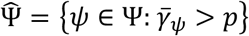, the subset of those branches tested that are inferred to be potentially antigenically significant by having inclusion probability above some threshold, *p*. In previous work these phylogenetic variables were selected using a random restart hill-climbing algorithm that optimised Akaike information criterion (AIC) to reduce computation cost (12,13). However, we have subsequently shown that variable selection using these binary masks is a superior strategy (18).

### Incorporating amino acid substitutions

Next, terms were introduced to explicitly attribute antigenic differences between viruses to specific amino acid changes. The shared evolutionary history of the viruses has the potential to facilitate false statistical support for substitutions due to repeated measurements. To control for the evolutionary relationship between viruses and reduce this risk, 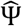, the subset of phylogenetic terms identified as explaining antigenic variation using **Eq 3**, were retained in the new model. Branch variables were ranked by their inclusion probability 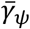 and p, the threshold inclusion probability, was chosen so that the the proportion of branches from Ψ that were carried forward in 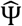 was 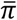, the posterior mean inclusion probability of a branch.

The subsequent model then included all of these terms (to control for the shared evolutionary history of the viruses) together with terms representing amino acid positions:

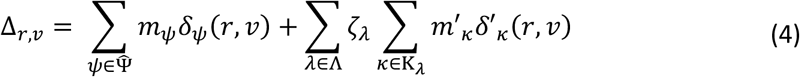

Here, each amino acid position within the set of non-conserved positions Λ is indexed by *λ*, while all observed amino acid substitutions at each position *λ* are in turn indexed by *κ*. Each position was associated with a binary mask variable, *ζ_λ_*, that takes the value zero or one, as above, determining whether substitutions at position *λ* contribute to variation in titres. A precomputed indicator variable, *δ*′*_k_*(*r, v*), indicates whether or not each specific amino acid difference separates the reference and test viruses. Each antigenic effect, *m*′*_κ_*, represents the antigenic effect of a specific substitution.

Using the model described in **Eq 4**, antigenic effects of alternative substitutions at the same amino acid position are independent. An alternative model where the antigenic effects of different substitutions at the same position are linked is also explored:

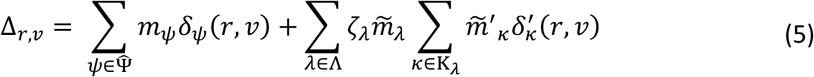

The key difference here from **Eq 4** is that the antigenic effect of a substitution is partitioned into a positionspecific component, 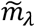, and a substitution-specific component, 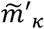. The position-specific component is shared by every substitution observed at an amino acid position. This reflects an expectation that a range of substitutions at more antigenically important positions will tend to have greater antigenic impacts than at other positions. The substitution-specific component, on the other hand, allows for variability in the antigenic impact of alternative substitutions at the same position.

### Incorporating structural information

Eqs 4 and 5, above, describe how variation in titres was attributed to antigenic effects of amino acid substitutions in what we term ‘structurally-naïve’ models. We now incorporate information from analysis of 3-D protein structure into the above models by using it to inform the prior probability that substitutions at an amino acid position are involved in antigenic evolution (by influencing the binary mask variable associated with each position). We implement this approach using two structural features: proximity to the receptor-binding site (RBS), and a predicted epitope score derived from a tool used to predict conformational epitopes from tertiary protein structure (see Materials and Methods). However, this approach is not limited to the use of these features and could be adapted to either a single or more than two structural features. Here, we refer to the two structural features as *F*_1_ and *F*_2_.

In Eqs 4 and 5 (the structurally-naïve models), the binary mask terms associated with each amino acid position, *ζ_λ_*, share a common prior distribution and therefore are equally likely to be included in the model prior to observation of the data. Here, the structurally-aware version of the models described above retain the structure described in Eqs 4 and 5. The only difference is that the binary mask variable 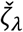 is redefined so that the probability of each position being selected may be influenced by structural features *F*_1,*λ*_ and *F*_2,*λ*_:

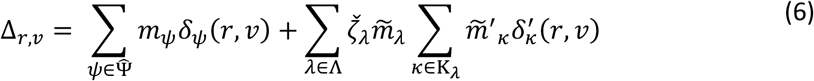

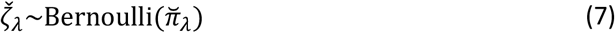

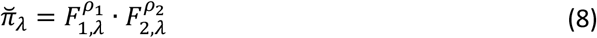

For each position, the outcome of Bernoulli trial which determines the value of the binary mask variable now depends on both the data and a position-specific, structure-informed prior probability of antigenic importance 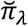. As described in **Eq 8**, the probability term 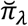 is computed directly from the structural features associated with position *λ* (which are scaled between zero and one) and the power terms *ρ*_1_ and *ρ*_2_, the values of which are fitted to the data. A tendency across all positions for those with higher values for the structural features to be involved in antigenic change will result in higher estimates for *ρ*_1_ and *ρ*_2_. Conversely, if the data do not support a relationship between a structural feature and antigenicity, the associated *ρ* term will tend towards zero.

## Results

To model the genetic basis of antigenic differences between influenza viruses, we first compiled antigenic (HI) and HA genetic sequence data from A(H3N2) viruses isolated during the period 1990-2014, and used the sequence data to construct a phylogeny describing the evolutionary relationships between viruses. Next, we mapped variation in HI titres to branches of the phylogenetic tree – BSSVS was used to identify branches of the phylogeny representing amino acid substitutions causing antigenic change and to quantify the associated degree of antigenic change. Specifically, we calculated the proportion of MCMC samples in which a phylogenetic variable from **Eq 3** is selected (where the associated binary mask variable, *γ_ψ_* = 1), which represents confidence in the selected variable, and we also recorded a quantitative estimate of the antigenic change, *m_ψ_* (in log_2_ HI units), associated with the branch.

The contributions of each branch to antigenic evolution are shown in the phylogenetic tree in **Fig 1a** where branch lengths indicate the posterior mean value of *γ_ψ_m_ψ_* in **Eq 3**. In this visualisation, the horizontal dimension expresses antigenic change. Consequently, an antigenically homogenous clade will appear as a flat vertical line, regardless of the amount of molecular evolution that has occurred within it or the time spanned. The mean number of branches included in the model in an individual step of the MCMC was 64 (95% HPD, 55-75). Branches in the trunk lineage were more likely to be included in the model explaining variation in HI titres, compared with branches in the rest of the tree (odds-ratio 5.0; 95% CI, 3.4-7.4). The histogram in **Fig 1b** illustrates a highly right-skewed distribution of antigenic weights assigned to branches with relatively few antigenic events of more than 1 log_2_ HI units. While the histogram shows branches with larger effects to be found in both the trunk and side branches, the rate of antigenic change was found to be considerably higher in the trunk. Rates of antigenic evolution in the trunk and side branches were calculated by summing antigenic weights (*γ_ψ_m_ψ_*) and dividing by the sum of branch lengths measured in years of evolutionary time estimated using a molecular clock analysis implemented in BEAST (20,21). The rate of antigenic drift in the trunk lineage was estimated to be 0.73 log_2_ HI units per year (95% HPD, 0.67-0.78), compared with 0.05 log_2_ HI units per year in the rest of the tree (95% HPD, 0.04-0.06). Cumulative antigenic distance from the root was calculated across the phylogenetic tree by summing antigenic effects (*γ_ψ_m_ψ_*) across each branch in a path between the root and every internal node and tip. This cumulative antigenic distance is represented in the horizontal dimension in the tree in **Fig 1a**. Linear regression of cumulative antigenic distance for each node and cumulative branch lengths estimated a rate of antigenic drift of 0.72 log_2_ HI units per year (**Fig 1c**). This aligns very closely with a figure of 0.71 previously estimated for a dataset of A(H3N2) viruses from the period 1985-2015 using a comparable approach (8).

**Figure 1.**
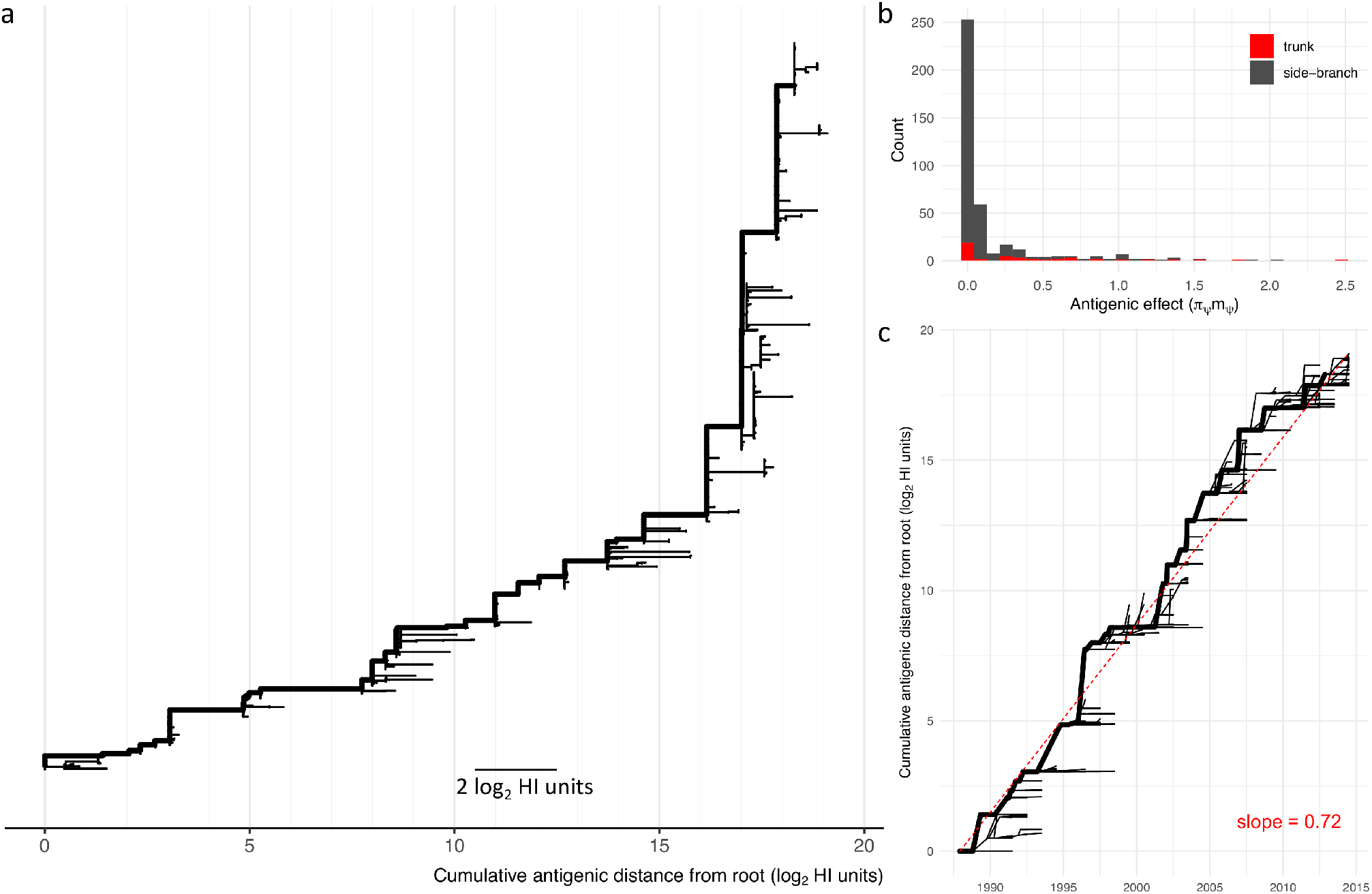
Antigenic evolution mapped to HA phylogeny. Antigenic change, as expressed in HI titres, mapped to branches of the HA1 phylogeny. The antigenic effect (log_2_ HI units) for each branch estimated as the average drop in titre when virus and antiserum separated by the branch are tested together. (**a**) HA phylogeny with branch lengths scaled to show antigenic effects (γ_ψ_m_ψ_). The x-axis shows cumulative antigenic distance from the root. The trunk lineage is shown as a thick line. (**b**) Histogram showing antigenic effects (γ_ψ_m_ψ_) estimated for each branch in the trunk lineage (red) and in the side branches (grey). (**c**) Phylogeny is plotted to show time on the x-axis (years) and cumulative antigenic distance from the root on the y-axis (log_2_ HI units). The trunk lineage is shown as a thick line. Dashed red line indicates the linear regression between time since the root and antigenic distance from the root for each node in the phylogeny (slope = 0.72).

### Genetic determinants of antigenic change

Next, amino acid substitutions were tested as predictors of reduced HI titres. Antigenic weights were estimated for each substitution with binary mask variables (*ζ_λ_*) dictating whether substitutions at a particular position were included in the genetic model of variation in titres. Based on the HI data, for each variable position the posterior mean value of *ζ_λ_*, or the posterior inclusion probability, is the inferred probability that substitutions at position *λ* have contributed to antigenic evolution. To assess how well each model did in terms of attributing antigenic changes to amino acid substitutions, we examined posterior inclusion probabilities associated with amino acid positions in the context of antigenic sites defined in the literature. While the important antigenic areas of HA are known, definitions of the constituents and boundaries of the antigenic sites vary, making a binary in-or-out classification to assess model selection problematic. For this reason, we considered the distance in 3-D space from each residue to these sites rather than a binary in-or-out classification. These distances were calculated between alpha carbons, therefore were relatively insensitive to the changes in protein structure occurring during evolution. For example, distances calculated using the HA structures of A/Aichi/2/68 and A/Brisbane/10/2007 were highly correlated (*R*^2^ >0.99, **Fig S1a**), despite the two viruses being separated by 39 years of evolution. At each step of the MCMC, the mean distance to an antigenic site, averaged across the set of positions selected by the model (*ζ_λ_* = 1) at that step, was calculated. The mean distance to antigenic site averaged over the MCMC chain was used to evaluate model performance with lower values indicating higher ability to correctly attribute antigenic variation to causative substitutions. Initial comparison of structurally-naïve models showed that a model assuming a link between the antigenic impact of subsitutions occurring at the same HA position (antigenic relationships, Δ*_r,v_*, modelled using **Eq 5**) outperformed a simpler model that assumed no such link (Δ*_r,v_* modelled using **Eq 4**) as indicated by a lower mean distance to antigenic sites of amino acid residues inferred to have substitutions explaining antigenic differences (Table 2). Using a single effect size (*m*′*_κ_* in **Eq 4** or 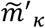 in **Eq 5**) for forward and reverse substitutions (symmetric substitution effects) also led to better model performance compared with estimating two effect sizes (asymmetric substitution).

**Table 2.** Model performance evaluated by distance from antigenic sites of positions implicated in antigenic evolution, averaged across posterior samples.

| Model | Symmetric substitution effects | Mean distance to antigenic site (Å) |
| --- | --- | --- |
| Structurally-naïve model - substitution model with independent effect sizes (**Eq 4**) | Yes | 11.80 |
|  | No | 13.28 |
| Structurally-naïve model - substitution model with effect sizes linked to position (**Eq 5**) | Yes | 6.86 |
|  | No | 11.39 |
| Structurally-aware model – structure influencing inclusion probabilities for positions and effect sizes for substitutions linked to position (**Eq 6**) | Yes | 2.55 |
|  | No | 3.05 |

For the best performing structurally-naïve model (described by **Eq 5**), inclusion probabilities for each position with substitutions present in the dataset are indicated in **Fig 2a**, with vertical bars showing the locations of antigenic sites (22). In **Fig 2b**, residues on a structural model of HA are shown coloured by inclusion probabilities and the locations of antigenic sites are shown to the right. Six HA positions were identified with very high confidence, each being associated with an inclusion probability of at least 0.95: positions 131, 135, 145 belonging to antigenic site A, 157 and 189 to antigenic site B, and 223 located on an exposed loop on the boundary of the RBS. Using the posterior mean value of the probability of inclusion, 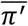, to determine the fraction of sites included results in 25 HA positions being selected. Of these 25 positions, six belong to antigenic site A (126, 131, 135, 137, 144 and 145), five to site B (157, 158, 159, 160 and 189), one to site C (53), two to site D (121 and 173) and none to site E. In **Fig 2c**, each HA position tested is positioned by its distance from the closest antigenic site and its posterior inclusion probability. When ranked by inclusion probability, the top 16 positions fall within 8Å of the core antigenic site limits, and only two positions above this threshold (183 at 9.0Å and 296 at 16.7Å) were included (with a probability greater than the threshold of 0.33). However, **Fig 2c** shows substitutions at some more distant positions were being included in the model in a non-negligible proportion of MCMC samples (positions 4 at 75.7Å and 14 at 52.3Å were associated with inclusion probabilities of 0.23 and 0.22 respectively). Both of these positions are in the stalk domain distant from the antigenic sites around the RBS and therefore it is deemed very unlikely that substitutions at these positions have contributed to antigenic evolution as assessed in HI assays. The occasional inclusion of parameters associated with positions such as these inevitably disturbs estimates for other parameters within the model.

**Figure 2.**
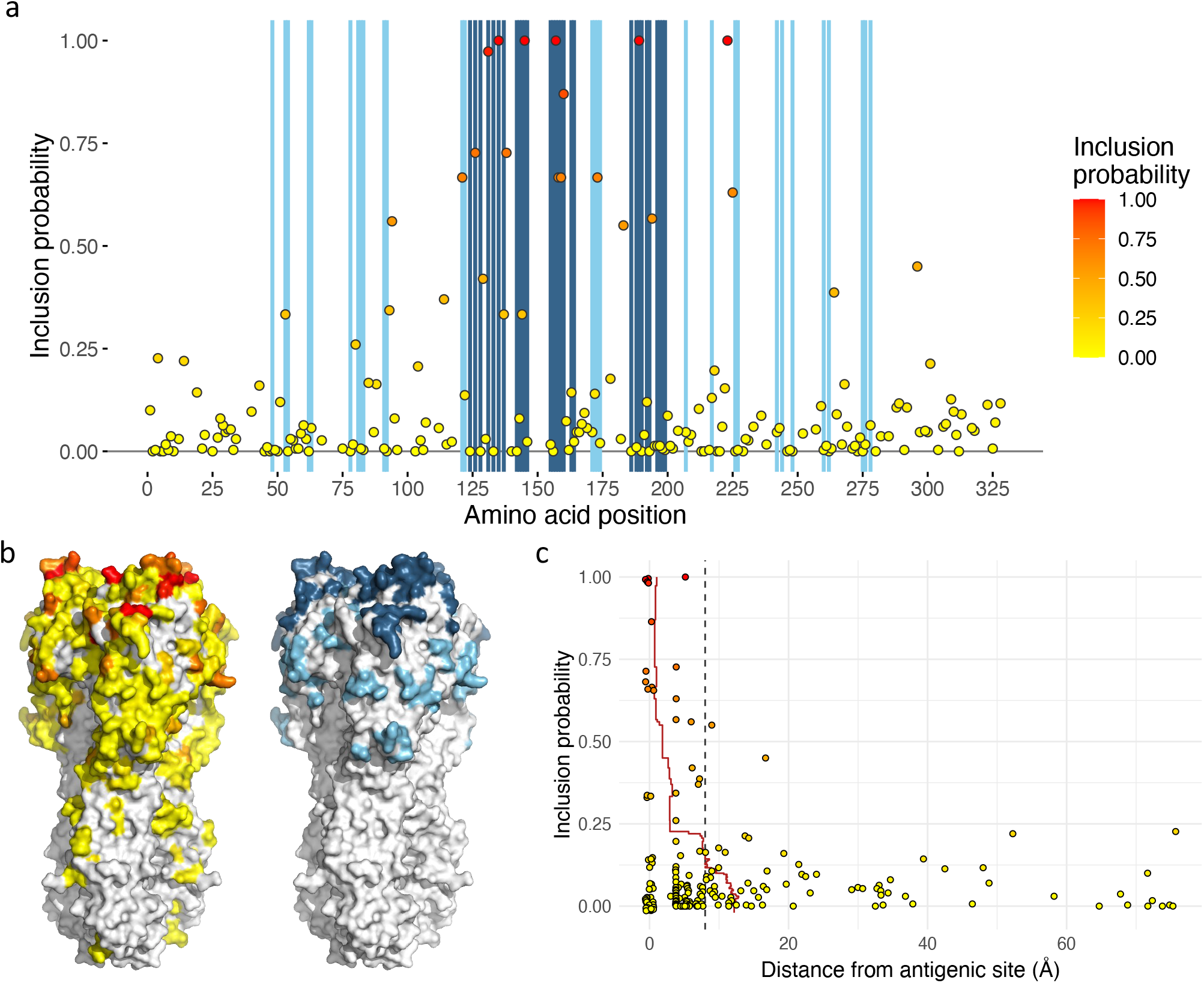
The role of HA positions in antigenic evolution estimated using a structurally naïve model. (**a**) Each point represents the posterior inclusion probability for a variable HA1 position. Blue vertical shading indicates positions on the x-axis that are described in antigenic sites A and B (dark blue) and antigenic sites C, D, and E (light blue). (**b**) Amino acid positions on the surface model of a HA structure are coloured: to the left, by inclusion probability following the colour scheme in **a**; to the right, antigenic sites A and B are shown in dark blue and sites C, D and E in light blue. (**c**) Posterior inclusion probability for each variable amino acid position is plotted against distance from the closest residue in a defined antigenic site. Points at 0Å on the x-axis, representing residues in described antigenic sites, have been adjusted in both dimensions to allow visualisation of overlapping points. A red line indicates the mean distance of residues selected with inclusion probability equal to or higher than that on the y-axis. A dashed vertical line at 8Å marks a threshold at which a position is approximately two residues away from an amino acid in a described antigenic site.

### Incorporating structural data

To investigate whether incorporating data on protein structure could assist the accuracy with which variables explaining antigenic differences could be identified, the locations of residues within the HA structure were used to influence the inclusion probabilities estimated for each position. Two structural features were considered, the distance from the RBS and a structure-based epitope score which estimates how accessible areas of a protein are to an antibody. For each residue, its distance to the RBS was calculated as minimum distance in 3-D space of its alpha carbon to the closest alpha carbon of a RBS residue. A structure-based epitope score was calculated for each HA residue using BEpro (23) and reflects how exposed and accessible for antibody binding, the region centred on each residue is. Comparison of these two measures calculated across solved HA structures from three A(H3N2) viruses isolated during the period covered by the HI dataset (A/Finland/486/2004, A/Hong Kong/4443/2005 and A/Brisbane/10/2007) and an earlier, evolutionary founder virus from the 1968 pandemic (A/Aichi/2/68) indicated that changes to protein structure during virus evolution did not greatly affect these measures. The variances in measurements made between different HA structures were small indicating that structural information used is not unduly influenced by the choice of HA structure. For example, comparison of RBS distances and epitope scores for the HAs of A/Aichi/2/68 and A/Brisbane/10/2007 were highly correlated (*R*^2^ >0.99 and 0.94 respectively, **Fig S1b-c**). Therefore, mean distance from the RBS and epitope scores averaged over the four structures (**Fig 3**) were considered suitable for use in modelling.

**Figure 3.**
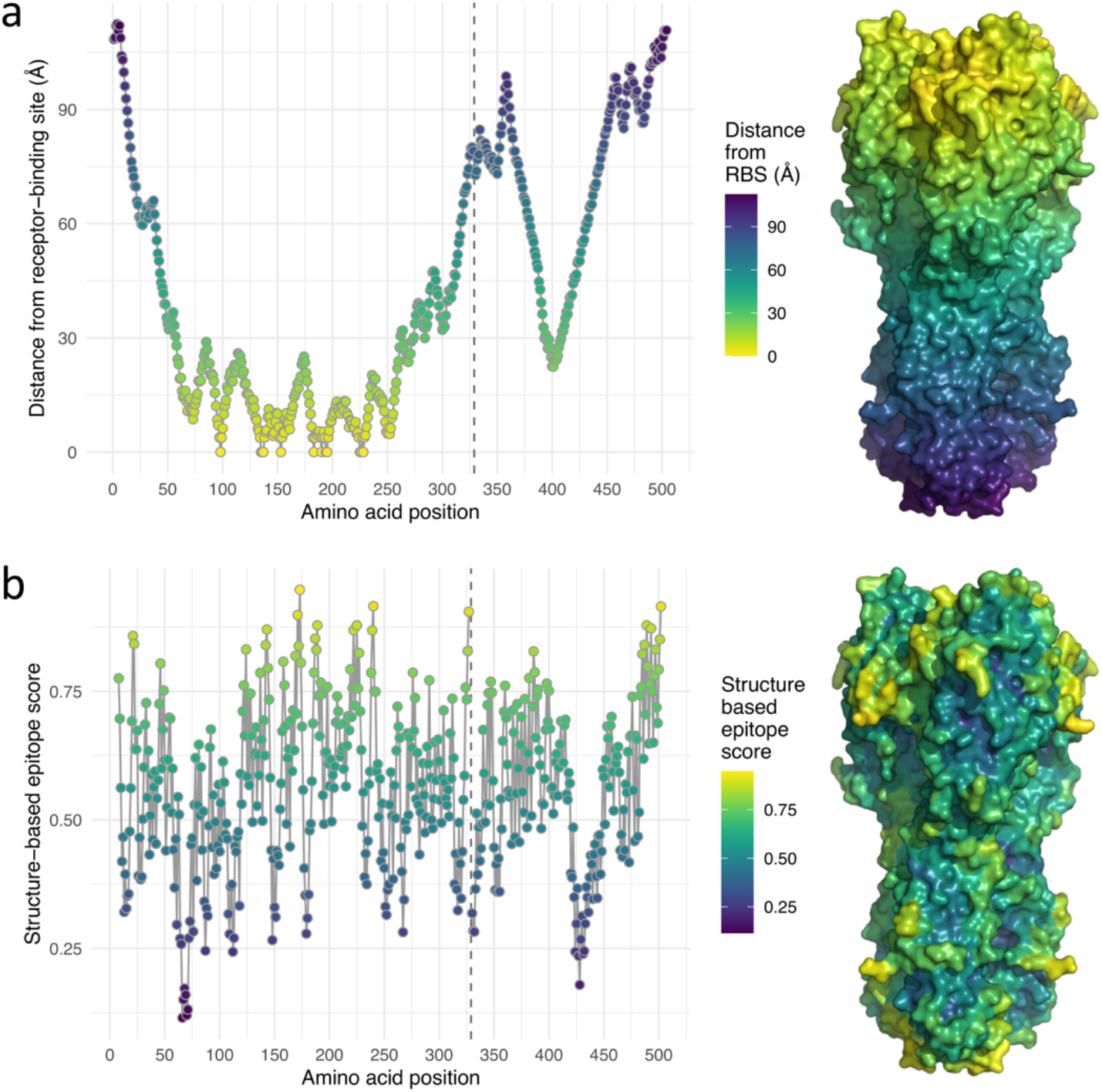
Structural features of the influenza A(H3N2) HA. (**a**) The distance of each HA residue to the closest of the residues comprising the RBS. To the right, a surface representation of the HA structure is shown coloured according to the distance key. (**b**) The structure-based epitope score for each HA residue was calculated using BEpro (23). To the right, a surface representation of the HA structure is shown coloured according to the epitope score key. In each plot, a vertical dashed line at position 329 indicates the boundary between HA1 and HA2.

The structural measurements plotted in **Fig 3** were allowed to influence the selection of amino acid substitutions inferred to explain variation in the antigenic component of HI titres, Δ*_r,v_*, according to **Eqs 6–8**. Structural information was used to estimate a structure-informed probability 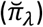 for each HA position, with the parameters *ρ*_1_ and *ρ*_2_ determining the importance of the proximity to the RBS (denoted *F*_1,*λ*_) and the structure-based epitope score (denoted *F*_2,*λ*_) of each position, according to **Eq 8**. In **Fig 4a** each HA residue is positioned in a 2-D space according to these two measures, while the colouring shows the structure informed probability of antigenic importance calculated from posterior mean values for the *ρ* parameters according to **Eq 8**. **Fig 4a** shows that neither the proximity to the RBS nor the structure-based epitope score fully dominated the determination of the probability, however **Fig 4b** shows the *ρ* parameter associated with RBS proximity (mean, 4.43; 95% HPD, 1.60-8.61) was higher than that associated with the epitope score (mean, 2.40; 95% HPD, 0.93-3.73), indicating proximity to the RBS to be particularly useful as a predictor, given both structure features were rescaled between zero and one for modelling. As expected, the structure-based probability does not fully determine whether or not a position is included in the model but guides variable selection, according to **Eq 7**, when correlations between patterns of substitution obscure relationships between antigenic change and causative substitutions. The correlation between the structure-based probability, 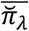, and the posterior inclusion probability, 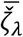, has a slope of 1.07 as shown in **Fig 4c**. Variation around the slope is expected as other factors such as the side-chain properties of the actual amino acid substitutions must contribute to determine whether positions influence antigenicity, however the slope of close to 1 indicates that the relationship between structure-informed probability and posterior inclusion probability is fitted correctly. In **Figs 4d-e**, HA positions are again placed according to structure-based epitope scores and distance from the RBS. Each non-conserved HA position is coloured by its posterior inclusion probability, the confidence that substitutions at the position have impacted antigenic evolution, in models fitted without (**Fig 4d**) and with structural data (**Fig 4e**). The effect of structural information guiding variable selection is clear with residues positioned in the bottom right corner of the scatterplot for the structurally-aware model tending to be associated with higher inclusion probabilities, which corresponds to a greater number of red residues in surface-exposed areas near to the RBS in **Fig 4e** compared with **Fig 4d**. Notably, a higher proportion of positions with an inclusion probability of one are identified using the structurally-aware model (**Fig 4f** compared with **Fig 2a**).

**Figure 4.**
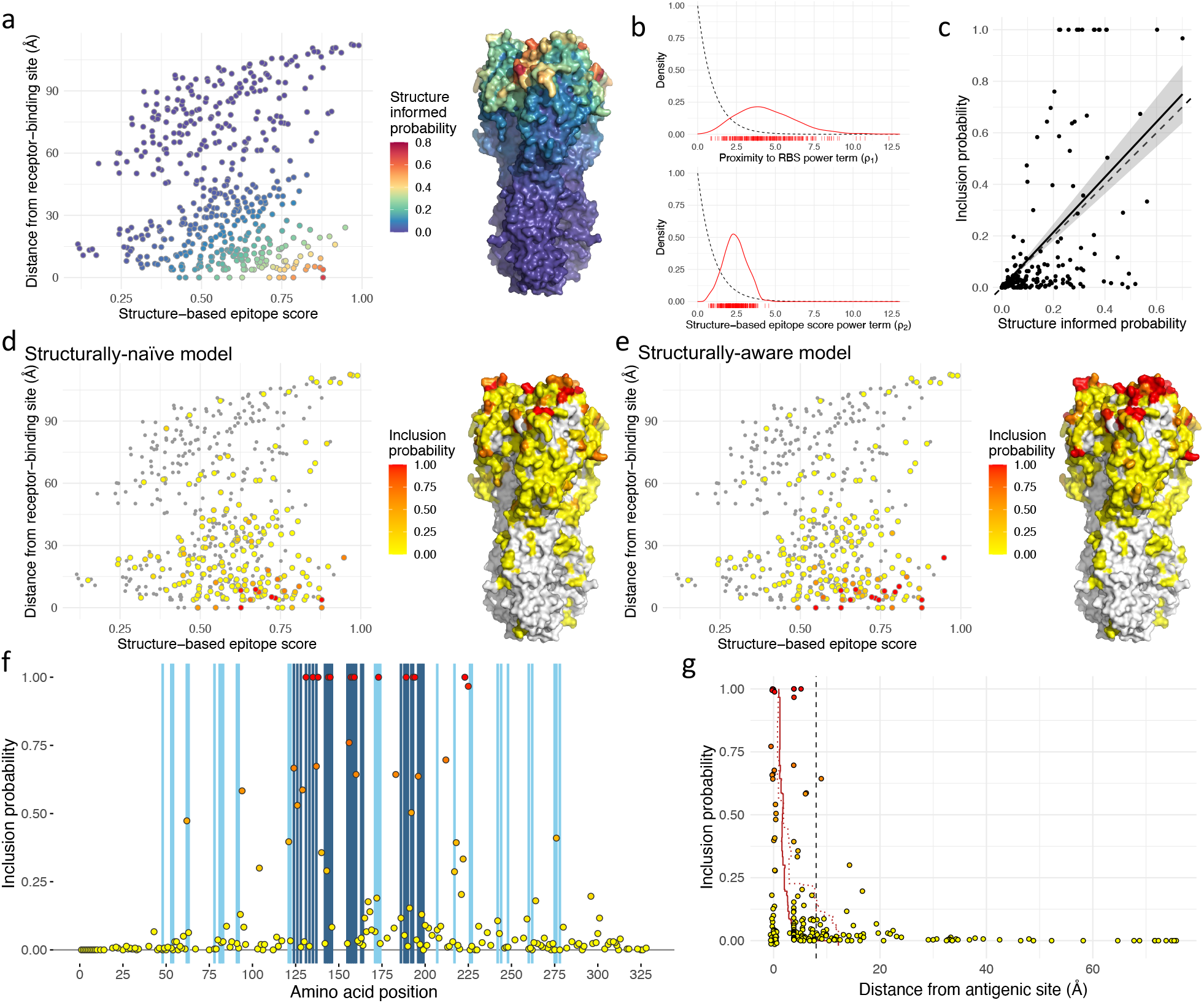
Use of structural data to guide variable selection. Plots summarise variable selection using a model where posterior inclusion probability that substitutions at an HA position affect antigenicity is impacted by a structure-informed probability that depends on a structure-based epitope score and distance from the RBS (**a**). HA residues are positioned according to a structure-based epitope score and distance from the RBS. Colour indicates the structure-informed probability of antigenic importance. To the right, structure informed probability is shown on the HA structure. (**b**) Posterior distributions for power terms that link proximity to the RBS (ρ_1_, top) and structure-based epitope score (ρ_2_, bottom) for each HA position to a structure-informed probability for the position, 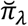, according to **Eq 8**. Individual values sampled from the posterior distribution are shown below the x-axis. Prior distributions for these parameters, defined as Gamma(1, 1), are shown as dashed black lines (**c**) Scatterplot showing the relationship between the structure-informed probability and the posterior inclusion probability. The solid black line has a slope of 1.07 alongside a dashed line of slope 1, with standard error from the linear model indicated in grey. (**d-e**) HA residues are positioned according to structure-based epitope score and distance from the RBS, identical to **a**. The colour scheme indicates posterior inclusion probabilities estimated using a structurally-naïve (**d**) and structurally-aware (**e**) models. Positions without substitutions are shown in grey. To the right of each scatterplot, inclusion probabilities are shown on the HA structure. (**e**) Each point represents the posterior inclusion probability for a variable HA1 position. Blue vertical shading indicates positions on the x-axis that are described in antigenic sites A and B (dark blue) and antigenic sites C, D, and E (light blue). (**g**) Inclusion probability for each variable amino acid position is plotted against distance from the closest residue in a described antigenic site. Points at 0 Å on the x-axis representing residues in described antigenic sites, have been adjusted in both dimensions to aid visualisation of overlapping points. A red line indicates the mean distance of residues selected with inclusion probability equal to or higher than that on the y-axis while a dotted red line shows the corresponding line from a structurally-naïve model shown in **Fig 2c**. A dashed vertical line at 8Å marks a threshold at which a position is approximately two residues away from an amino acid in the described antigenic site.

Incorporating structural information into variable selection reduced uncertainty in model identification. Using the structurally-aware model, 144 of 199 (72.4%) variable HA positions were either included in the model or excluded in at least 95% of MCMC samples (inclusion probability >0.95 for 14 positions and <0.05 for 130, **Table S1**), while for the structurally-naïve model the corresponding number was only 119 (59.8%) (>0.95 for 6 positions and <0.05 for 113, **Table S1**). In **Fig 4f**, posterior probabilities are shown for each HA position where substitutions were present with blue shading used to indicate antigenic sites. This plot shows that the majority of selected positions either belonged to defined antigenic sites or were very close to one of them in primary amino acid sequence. Of the 14 positions associated with an inclusion probability of at least 0.95, four were in antigenic site A (positions 131, 135, 144 and 145), five in site B (157, 158, 159, 189 and 193) and one in site D (173), while the others were either defined as belonging to the RBS (194 and 225) or were located close to the RBS (138 and 223). Incorporating structure into variable selection resulted in greater accuracy as quantified by the distances of amino acid residues from defined antigenic sites (**Fig 4g**, **Table 2**). Each of the 18 residues with the highest inclusion probability belonged to defined antigenic sites or were within 8Å of these and only position 183, at 8.9Å, was further away and associated with an inclusion probability of >0.5. Of all positions associated with an inclusion probability of >0.05, position 43, 19.3Å away, with an inclusion probability of 0.08, was the most distant. For comparison, 19 positions further than 19.3Å from antigenic sites were associated with inclusion probabilities of at least 0.05 using the structurally-naïve model (**Fig 2c**), up to a maximum of 75.7Å (position 4, inclusion probability of 0.23).

### Bayesian model averaging delivers accurate prediction of HI titres

The full dataset was comprised of 38,757 titres which included 3,477 different virus and reference antiserum combinations. These combinations were measured with 1,737 different viruses (including reference viruses) and antisera raised to 151 reference viruses. To assess the predictive performance of genetic models of antigenic phenotype, a range of model variants were tested for their capacity to recover HI titres under two prediction schemes. Firstly, to assess the capacity of models to predict unobserved antigenic relationships, 10% of virus and reference antiserum combinations (348 combinations), were randomly selected 100 times, and all titres associated with those combinations were removed to act as test data. Models were trained using the remaining 90% of virus and reference antiserum combinations and tested for their ability to accurately predict the removed titres. Secondly, to evaluate model capacity to predict titres of uncharacterised viruses, 10% of viruses (174 viruses) were randomly selected 100 times, and all titres for these viruses measured using any antiserum were removed to act as test data with data for the remaining 90% of viruses used to train models. Under both prediction schemes, the models (**Eqs 3–6**) included indicator variables to determine whether a genetic variable contributed to differences in titres, in common with the previous sections. As these indicator variables are present, the combination of HA positions at which substitutions contribute to predictions may vary at each step of the MCMC. This therefore constitutes prediction by model averaging rather than prediction conditioned on a single best model, accounting for uncertainty in the identification of the substitutions that cause antigenic differences.

Employing the first scheme, titres were predicted initially with the antigenic component of titres, Δ*_r,v_*, modelled using the phylogenetic model described by **Eq 3**, with information on the antigenic changes associated with branches of the phylogeny (*γ_ψ_m_ψ_*) estimated using the training data only. This model provides a reasonable approximation of antigenic relationships, allowing titres to be predicted for unknown virus and reference strain combinations under cross-validation with a mean absolute error (MAE) of 0.60 log_2_ HI units (root-mean-square deviation RMSD = 0.83, **Table 3**, scheme 1). However, as the antigenic weights (*γ_ψ_m_ψ_*) associated with branches are purely additive, the tree model fails to account for the antigenic consequences of phenomena such as reverse substitutions or the presence of the same substitution in multiple branches.

**Table 3.** Accuracy of genetic models in predicting antigenic phenotype. Measures of the difference between predicted and observed HI titres and the percentage of errors within 1 or 2 log_2_ HI units.

| Model (Equation) | Symmetric substitution effects | Prediction scheme |  |  |  |  |  |
| --- | --- | --- | --- | --- | --- | --- | --- |
|  |  | 1. Test-reference virus pairs |  |  | 2. Test viruses |  |  |
|  |  | MAE | RMSD | % <1 (<2)<sup>1</sup> | MAE | RMSD | % <1 (<2)<sup>1</sup> |
| Phylogenetic model (**Eq 3**) | n/a | 0.60 | 0.83 | 83 (94) | 1.03 | 1.14 | 64 (87) |
| Structurally-naïve model - substitution model with independent effect sizes (**Eq 4**) | Yes | 0.53 | 0.69 | 86 (99) | 0.80 | 1.04 | 69 (94) |
|  | No | 0.53 | 0.69 | 86 (99) | 0.81 | 1.05 | 69 (93) |
| Structurally-naïve model - substitution model with effect sizes linked to position (**Eq 5**) | Yes | 0.52 | 0.69 | 87 (99) | 0.80 | 1.04 | 69 (94) |
|  | No | 0.52 | 0.69 | 87 (99) | 0.79 | 1.03 | 70 (94) |
| Structurally-aware model – structure influencing inclusion probabilities for positions and effect sizes for substitutions linked to position (**Eq 6**) | Yes | 0.52 | 0.69 | 87 (99) | 0.77 | 1.01 | 71 (94) |
|  | No | 0.52 | 0.69 | 87 (99) | 0.77 | 1.02 | 71 (94) |

MAE, mean absolute error; RMSD, root-mean-square deviation; n/a = not applicable. <sup>1</sup>The percentage of predicted titres within 1 (or 2) $\log_2$ HI units of the true underlying titre.

To account for these non-additive antigenic events requires terms that explicitly describe the presence or absence of amino acid substitutions (Δ*_r,v_* modelled using **Eq 4, 5** or **6**). Using a model with independently estimated effect sizes for every substitution at each position (**Eq 4**) resulted in more accurate predictions with a MAE of 0.53 log_2_ HI units and RMSD of 0.69 (**Table 3**, scheme 1). Having the effects of substitutions at the same position linked (**Eq 5**) resulted in a marginal improvement in predictions (MAE = 0.52 and RMSD = 0.68) (**Table 3**, scheme 1). Improved predictions are expected under these substitution models, as they can account for convergent substitutions and reversions of substitutions. Using the structurally-aware model of antigenic relationships (**Eq 6**) resulted in similar prediction accuracy when predicting missing test-reference virus combinations (MAE = 0.52 and RMSD = 0.69, **Table 3**, scheme 1). The accuracy of these predictions are comparable to existing models for prediction of A(H3N2) HI titres (8). This indicates that our efforts to increase the stringency of variable selection, rather than prioritising the selection of the maximally predictive set of genetic terms, do not hamper the predictive ability of our approach. Under the best performing genetic model, 87% of predictions were made within 1 log_2_ HI unit of the observed titre and 99% within 2 log_2_ HI units (**Table 3**).

Under the second scheme, titres were predicted for 10% of viruses that had all titres removed to act as test data. The accuracy of all models was reduced under this prediction scheme (**Table 3**, scheme 2) with a MAE of only 1.13 log_2_ HI units (RMSD = 1.14) using the phylogenetic model. Predicting titres for viruses that are entirely absent from the training data is more challenging, in part, as it is not possible to estimate a virus avidity parameter (*A_v_* in **Eq 2**), as recognised previously (8,12). The accuracy of predictions was improved using the substitution-based models (MAE 0.77-0.81 log_2_ HI units and RMSD = 1.01-1.05). Interestingly, the structurally-aware model is slightly better performing here when predicting for cross-validation test datasets consisting of missing test viruses (0.77 compared with 0.79, **Table 3**, scheme 2). This indicates that more accurately attributing antigenic variation to the correct substitutions also offers an advantage when predicting antigenic relationships from HA sequences for viruses with no associated antigenic data.

In **Fig 5a**, titres predicted using scheme 1 are plotted against observed titres for the model with structural information (**Eq 6**). In **Fig 5b**, titres predicted using scheme 2 are plotted against the observed titres using the same model. **Fig 5a** shows a close relationship between predicted and observed titres while **Fig 5b** shows that in the absence of any information on the reactivity of a virus with any available antisera, there is a trend towards under-estimation of high observed titres. Such high titres tend to be associated with test viruses having higher than average titres across panels of antisera against which they are tested. Without information on these viruses (scheme 2), we observe a mean underestimation (predicted *v* observed) of - 0.16 (**Fig 5b**), which compares with a corresponding value of −0.06 for scheme 1 (**Fig 5a**).

**Figure 5.**
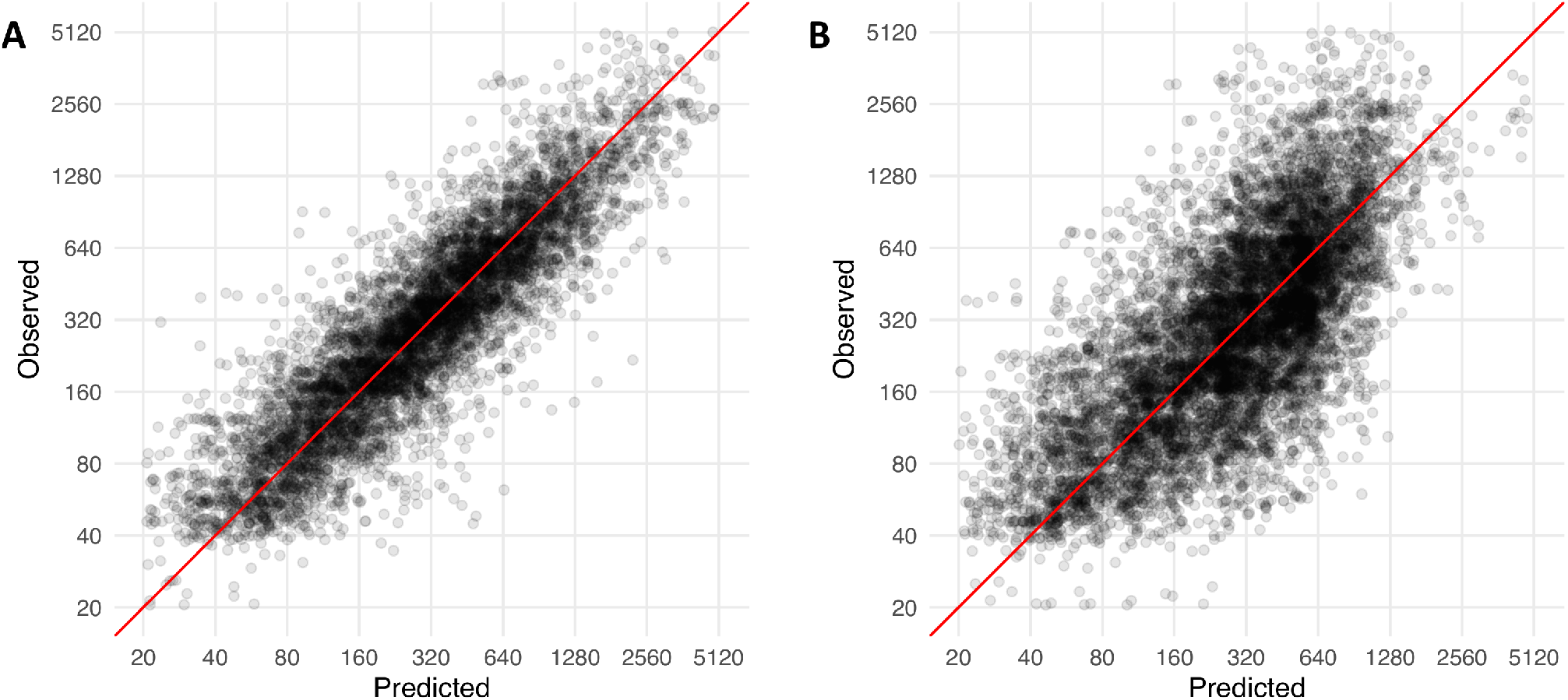
Sequence-based predictions of antigenic phenotype. Measured HI titres are plotted against predictions made under cross-validation procedures where test sets consist of randomly selected (**a**) 10% of test-reference virus pairs and (**b**) 10% of test viruses. Predictions are made using a structurally-aware model (**Eq 6**) in which structural features associated with the location of each residue in the HA protein influence the combination of genetic terms that contribute to predictions. Alternative substitutions at the same amino acid position vary in effect size though they are linked by a position-dependent component of the variable. Observed titres are the fitted titre for each reference strain and test virus, accounting for day-to-day variability in measured titres

## Discussion

Here, we describe an approach for identifying genetic changes that explain the antigenic drift of a rapidly evolving virus pathogen. The evolutionary process results in highly correlated genetic signals which can prove challenging for accurate mapping of genotype to antigenic phenotype. We describe a Bayesian approach whereby the causative amino acid changes are identified using BSSVS, performed alongside model fitting. Incorporating phylogenetic structure into the analysis favours amino acid positions where there are substitutions correlating with antigenic change at multiple points in the evolutionary history of the virus. The model additionally allows different substitutions at the same position to vary in antigenic impact and the option to link the antigenic effects of different substitutions occurring at the same position. Moreover, we show how information on the structural context of amino acid positions can guide the selection of antigenically relevant genetic variables for the HA of influenza A(H3N2) viruses, increasing the proportion of variables either included or excluded with high confidence (from 59.8% to 72.4%) and increasing the tendency for the model to attribute antigenic variation to substitutions at HA positions within or nearby recognised antigenic sites (reducing the mean distance across samples from 6.86 Å to 2.55 Å). Cross-validation demonstrates that this approach can additionally provide accurate predictions of antigenic phenotype from HA gene sequence.

The evolutionary process by which viruses are derived can result in multiple mutations occurring at the same (or almost the same) location in the phylogeny. When multiple amino acid substitutions result and these occur together at a point where antigenic change occurs, there is uncertainty as to which substitution(s) have caused the antigenic change. Information on the antigenic consequences of the same substitutions occurring in other branches of the phylogeny or other substitutions at the same amino acid positions can be used to resolve these ambiguities in some cases, but often uncertainty remains and it may be impossible to define a single definitive set of causative genetic changes by statistical means. A benefit of a Bayesian approach with indicator variables is is that this explicitly accounts for uncertainty in the identification of causative amino acid substitutions and reflects this uncertainty in variable selection. A further advantage of a Bayesian approach in this context is the opportunity to include prior information on the antigenicity of HA, here in the form of structural data; something we show can help to further resolve ambiguities in model selection. Model uncertainty was continued into cross-validation with predictions of masked titres made using Bayesian model averaging over several models weighted by the posterior probability of each model given the data, rather than a single model with a single set of causative amino acid substitutions.

We describe an approach to integrate data on proximity to the RBS and a pre-computed structure-based epitope score into priors for the probability that substitutions at a position impact antigenicity, an approach that could be easily extended to other proteins or structural features. In this analysis of the influenza HA, a prior expectation that antigenically important substitutions would tend to occur at HA positions both increased proximity to the RBS and higher epitope scores was implemented through choice of priors. However the model was free to minimise the influence of either parameter and this approach is not limited to situations in which prior knowledge of the directionality of a relationship exists. An extension of this work would be to account for the shielding of epitopes by covalently attached glycans and the changes in the accessibility of epitopes that occurs as a result of changes in glycosylation over time (24,25). Our analysis was focused on the HA protein as the HI assay measures antigenic variation in HA. However, the analysis could be extended to also include neuraminidase to test for an antigenic effect of substitutions in that glycoprotein which may influence VN assays. There is also scope to exploit the transfer of information from data-rich to data-starved virus subtypes. For example, the relationship between structural features and the role in antigenic evolution could be trained using a data-rich subtype such as A(H3N2); this could inform the prior distributions for the inclusion probabilities of structurally-aligned positions in any newly emerged subtype that would have fewer available data.

Having developed the genetic model of antigenicity to fit known data, its predictive power was evaluated by cross-validation. Notably, we tested our methods by removing 10% of samples from the full dataset and used the remaining 90% of the data as the training set. This was done in two ways. The first was to remove 10% of virus and antiserum combinations, leaving other antigenic data for a particular virus in the training dataset. In this scheme (scheme 1) we saw that we could estimate as much as 87% of titres within 1 log_2_ HI unit of the observed titre and 100% within 2 log_2_ HI units and **Fig 5a** shows accuracy across all values of observed titre. This would mean, for example, that if we knew some of the characteristics of a test virus we could accurately estimate its HI titre against a new reference antiserum, a vaccine virus antiserum or a candidate vaccine virus antiserum within a two-fold dilution in most cases. The accuracy of the prediction was improved by the inclusion of variables representing amino acid substitutions (**Eqs 4–6**) over the basal phylogenetic model (**Eq 3**) which, lacking terms representing specific amino acid differences, cannot recognise when clades separated on the tree are antigenically similar because they share substitutions occurring convergently in different branches. Predictions were not markedly enhanced by linking the magnitude of antigenic effects caused by different amino acid substitutions occurring at the same HA position (**Eq 5**), or by the inclusion of structural information to guide the identification of the substitutions (**Eq 6**).

In the second, more challenging, assessment of the methods we removed all antigenic data for a virus and predicted its titres against reference antisera. Here, predictions were somewhat less accurate though up to 71% of predicted titres were within 1 log_2_ HI unit of the observed titre and 94% within 2 log_2_ HI units. In this prediction scheme (scheme 2) shown in **Fig 5b**, it is clear that the model rarely predicts high HI titres (e.g. >1280) with no knowledge of whether the ‘missing’ test virus has an inherent ability to be more sensitive to specific or non-specific inhibition by antisera. This result would, in the absence of any established antigenic data, predict whether an unknown virus was not well recognised by an antiserum (recognition of the test virus by antisera raised against reference viruses at titres 4-fold lower than the titre with the homologous virus) or was poorly recognised by such antisera (recognition at titres >4-fold lower than the homologous titres). Again, the greatest enhancement was, as seen in scheme 1, due to the inclusion of specific amino acid substitutions over and above the basal phylogenetic model. However, in scheme 2, the accuracy of the predictions improved, first by inclusion of specific amino acid substitutions (**Eq 4**), then by linkage of antigenic weights of alternative substitutions at a position (**Eq 5**), and finally by including information on the proximity of residues to the RBS and structure-based epitope scores (**Eq 6**).

In summary, the ability to predict antigenic cross-reactivity of emerging influenza viruses, as measured by HI, while maximising identification of the causative amino acid substitutions provides important information with which to evaluate the epidemic potential of influenza virus variants. Using models parameterised using data collected in previous years can help to refine such techniques and we describe how structural data can be incorporated into model fitting. Incorporating other sources of prior information is an exciting area for further model development, for example alternative protein structural data could be tested as explaining variation in alternative assays used to characterise antigenic similarity of viruses. The benefits of being able to integrate such data types into modelling of evolution could be particularly powerful when performing analyses of emerging influenza viruses for which historic data are unavailable. The approach we describe, allowing detailed and accurate mapping of genotype to antigenic phenotype, should progress efforts to understand the genetic determinants of virus fitness and evolutionary trajectories of influenza viruses, importantly when surveillance is increasingly based on a ‘sequence-first’ approach. Moreover, this approach could also be adapted to proteins of other viruses such as the capsid proteins of foot-and-mouth disease virus and the spike protein of coronaviruses.

## Materials and Methods

### Influenza data

Influenza viruses were originally isolated from clinical specimens either by WHO National Influenza Centres or by the London-based WHO Collaborating Centre (CC). The antigenic dataset for A(H3N2) included 1737 viruses for which HI and HA1-encoding gene sequence data were generated at the CC. All HI data used were from assays carried out at the CC and were obtained using post-infection ferret antisera. The data associated with this study are available online (26). HA1 nucleotide sequences and collection dates were analysed to generate temporal phylogenies using BEAST v1.8.2 (27). Phylogenies were estimated using a variety of nucleotide substitution, demographic, and molecular clock models. A general time reversible model of nucleotide substitution with proportion of invariant sites and a gamma distribution with four categories describing among-site variation (GTR + / + Γ_4_) was determined to be the most suitable model by comparison of Bayes factors. The trunk lineage was defined from the root through the descendant node leading to the greater number of sampled viruses.

### Structural analysis

For each residue in the structure, the distance from the RBS was calculated as the minimum distance in 3-D space between the alpha carbon of that residue and the nearest alpha carbon of a residue in the RBS (positions 98, 135, 136, 153, 183, 190, 194, 195, 225, 226, and 228) (28). The distance of each HA residue to the nearest antigenic site constituent was calculated as the minimum distance in 3-D space from the residue’s alpha carbon to the nearest alpha carbon of a residue in the antigenic site A (positions 124, 126, 128, 131, 133, 135, 137, 142, 143, 144, 145, and 146), site B (155, 156, 157, 158, 159, 160, 163, 164, 186, 188, 189, 190, 192, 193, 196, 197, 198 and 199), site C (48, 53, 54, 275, 276, 278), site D (121, 122, 171, 172, 173, 174, 207, 217, 226, 227, 242, 244 and 248) and site E (62, 63, 78, 81, 82, 83, 91, 92, 260 and 262) (22,29,30). To determine structure-based epitope scores for each residue in the HA structure from tertiary structure, the program BEpro (23) was used to analyse structures in Protein data bank (PDB) format. These scores reflect side chain orientation and solvent accessibility calculated using half sphere exposure values at multiple distances and amino acid propensity scores. For each residue, both half sphere exposure measures and propensity scores depend on all atoms within 8-16Å of the target residue, with increased weighting towards nearer atoms. Due to this, scores for any given residue are relatively insensitive to the effects of single amino acid substitutions. The BEpro server was accessed at http://pepito.proteomics.ics.uci.edu/info.html. Structural features were calculated from the solved HA structures of three human A(H3N2) viruses isolated during the period covered by the HI dataset, A/Finland/486/2004 (PDB: 2YP2 (31)), A/Hong Kong/4443/2005 (PDB: 2YP7 (31)), and A/Brisbane/10/2007 (PDB: 6AOU (32)) and from an earlier virus isolated in 1968 (A/Aichi/2/68 PDB: 3HMG (33)).

### Model Implementation

The *Modelling Approach* section above describes the modelling process by which variation in HI titres was attributed to genetic differences between viruses while accounting for phylogenetic relationships and non-antigenic sources of variation in titres using a hierarchical model structure. The terms used in the models are described in **Table 1**. This section describes choices of prior and other implementation details. Models were fitted in JAGS v4.3.0 (34) using the R package runjags v2.0.4-6 (35). Post-infection ferret antisera were raised to a range of reference viruses (*r*) and HI titres measured for an individual antiserum and several viruses (*v*) (including the homologous titre to the corresponding reference virus *r* and a range of other viruses). The measured titre for antiserum against reference virus *r* and virus *v* on a given date *d, Y_r,v,d_* was modelled as an underlying titre for the combination of reference virus and test virus pair, *H_r,v_*, and an effect for date along with a variance term (**Eq 1**). To reflect a lack of prior information, the effect for date was implemented with a diffuse prior for the effect of date defined as a normal distribution where 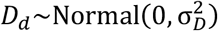 with the variance parameter 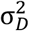 was drawn from an inverse gamma distribution such that 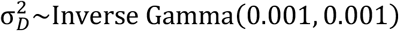. The variance term accounting for residual variation in measured titres was drawn from the prior distribution 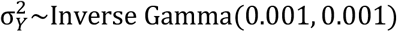. The underlying titre for each test virus and reference strain was modelled according to a general structure described by **Eq 2**. Priors for the estimated impact on titres associated with the use of antiserum raised using each reference virus were defined as 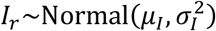. Here, the mean was given the prior 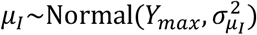 where *Y_max_* was the maximum observed titre and 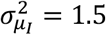 to maintain a value for the intercept proximal to the range of possible recorded titres without explicitly enforcing a particular value such as the maximum observed titre within the dataset and 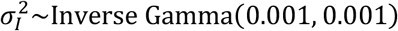. The prior for *μ_I_* was chosen to allow a fitted model with a high intercept from which lower titres were fitted by the subtraction of positive antigenic terms. Effects associated with the use of each test virus were associated with the prior 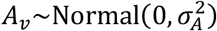 where 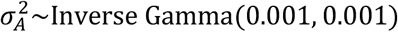. The variance term accounting for residual variation in underlying titres was drawn from the prior distribution 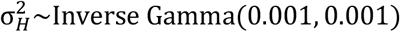.

The antigenic component of the model described in **Eq 3**, Δ*_r,v_*, estimates the drop in HI titres caused by the antigenic dissimilarity of reference strain *r* and test virus *v*. Δ*_r,v_* was modelled in several ways (**Eqs 3–6**) though was restricted to be positive throughout and was always subtracted from the other terms contributing to variation in titres (**Eq 3**). Using the combination of branches of the phylogenetic tree traversed when a path between them is drawn through the tree (**Eq 3**), each branch, *ψ*, of the phylogenetic tree, Ψ, was associated with a binary mask term, *γ_ψ_*, that determined whether the branch *ψ* was included in the model or not and an antigenic weight parameter, *m_ψ_*, representing the estimated drop in HI titres due to antigenic change mapping to the branch. These antigenic weights were required to be non-negative and the prior was defined as *m_ψ_*~ Gamma(2, 1). This choice of prior distribution discouraged the inclusion in the model of a high number of branches associated with very small antigenic weights, on the basis that effects very close to zero cannot be identified in HI assay data, thereby encouraging a more parsimonious model. Binary mask variables associated with each branch were drawn from a Bernoulli trial defined as *γ_ψ_*~Bernoulli(*π*) where *π* was given the prior *π*~Beta(2, 8) to favour a sparse model reflecting the expectation that a relatively low proportion of branches were expected to represent genetic differences influencing titres. The posterior mean value of *π* determined the proportion of branches used to account for phylogenetic structure when testing variables representing specific amino acid differences (Eqs 4, 5 and 6).

Next, terms were introduced to explicitly attribute antigenic differences between viruses expressed in HI assays to specific amino acid differences (**Eq 4**). These terms representing specific amino acid difference were tested in the presence of a subset of phylogenetic terms selected using **Eq 3**, 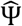. Each amino acid position was associated with a binary mask term, *ζ_λ_*, for which the prior was specified as *ζ_λ_*~Bernoulli(*π*′) where given the prior *π*′~Beta(2, 8) mirroring that used for the binary mask associated with phylogenetic terms above. When implementing **Eq 4**, priors for the effect sizes associated with substitutions were specified as *m*′_*κ*_~ Gamma(1, 1), which allows for antigenic effects close to zero - necessary as this allows for substitutions that do not affect antigenic cross-reactivity to occur at amino acid positions included in the model due to the presence of other antigenically important substitutions at the position. A similar model where substitutions occurring at the same amino acid position are linked in their effect size is described by **Eq 5**, which differs from **Eq 4** by having the antigenic effect of a substitution partitioned into position-specific and substitution-specific componenets, 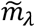 and 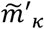 respectively. Priors for the position-specific antigenic parameter were specified as 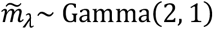, discouraging near-zero antigenic effects at the position level. **Eq 5** was implemented with the prior for specific amino acid substitutions specified as 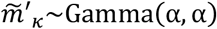 where *α*~Gamma(2, 1). This allows for a range of small and large antigenic impacts and, but the mean of this prior is equal to 1 regardless of the value of α. Therefore, for rarer substitutions (ones informed by very few titres) where the data make reliable estimation of an effect size unlikely, 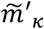 tends towards 1, so the antigenic impact of the substitution will be largely determined by the value estimated for the position-specific antigenic effect 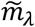.

Structural data associated with each amino acid position were used to influence the probability that substitutions at that position contribute to antigenic evolution as apparent in HI titres (**Eqs 7 and 8**). The binary mask term for each position, 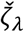, now depends on a position-specific structure-based probability of antigenic importance 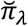. Two features derived from structural analysis of published H3 HA structures, as described above, were defined for each HA position: the distance from the RBS, *F*_1_, and a structure-based epitope score, *F*_2_ (see Structural analysis section). Each of these were re-scaled between zero and one prior to modelling so that higher values reflected high proximity to the RBS and high epitope scores respectively. Whether or not substitutions at an HA position contributed to variation in HI titres depended on the outcome of a Bernouilli trial where for each position, *λ*, a structure-informed probability 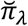 was determined as the product of *F*_1,*λ*_ and *F*_2,*λ*_. Higher values of each structural feature were expected to increase the probability that a position was antigenically important and so the parameters *ρ*1 and *ρ*2 were restricted to be positive and their priors were defined as *ρ*_1_~ Gamma(1, 1) and *ρ*_2_~ Gamma(1, 1).

### Cross-validation using Bayesian model averaging

Two cross-validation schemes were performed with the full dataset repeatedly divided into training and test datasets at random, criteria for each are described in the Results section. Under each scheme, measurements for antisera not present in the training data were excluded from the test dataset. Models described by **Eqs 3–6** were each fitted to the training data and used to predict HI titres for virus and antiserum combinations present in the test data. Predicted titres were compared with observed titres, with both MAE and RMSD calculated. Each error influences MAE in direct proportion to the absolute value of the error whereas RMSD placing more emphasis on penalisation of higher errors.

## Abbreviations

AIC: Akaike information criterion
BEAST: Bayesian evolutionary analysis sampling trees
BEpro: Discontinuous B-cell epitope prediction
BSSVS: Bayesian stochastic search variable selection
GISRS: Global Influenza Surveillance and Response System
GTR: generalised time-reversible
HA: haemagglutinin
HI: haemagglutination inhibition
HPD: highest posterior density
LASSO: least absolute shrinkage and selection operator
JAGS: Just Another Gibbs Sampler
MAE: mean absolute error
MCMC: Markov chain Monte Carlo
PDB: Protein Data Bank
RBS: receptor-binding site
RMSD: root-mean-square deviation
(e)SABRE: (extended) Sparse hierArchical Bayesian model for detecting Relevant antigenic sites in virus Evolution
VN: virus neutralisation
WHO: World Health Organisation

## Supplementary Material

**Figure S1.**
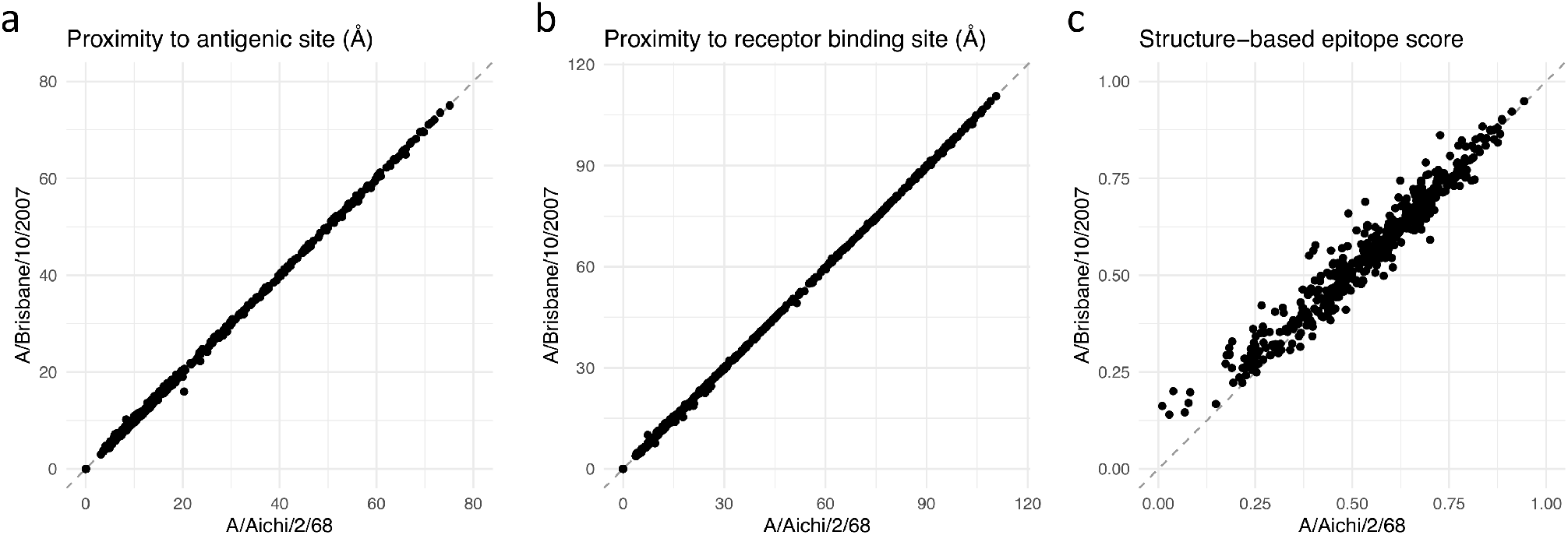
Correlation between structural features of A/Aichi/2/68 and A/Brisbane/10/2007 HA. (**a**) The correlation in the distance of the alpha carbon of each HA residue to the closest alpha carbon of a residue belonging to an described antigenic site. (**b**) The correlation in the distance of the alpha carbon of each HA residue to the closest alpha carbon of a residue belonging to the receptor-binding site. (**c**) The correlation in structure-based epitope scores estimated for each HA residue using the software BEpro.

**Table S1.** Model confidence in variable selection and distance of included or excluded HA positions to known antigenic sites.

|  |  | Number of positions<sup>a</sup> with given posterior inclusion probability and distance from antigenic site |  |  |  |
| --- | --- | --- | --- | --- | --- |
|  |  | Confidently included<br>(Inclusion prob >.95) |  | Confidently excluded<br>(Inclusion prob <.05) |  |
| Model | Symmetric substitution effects | N positions | Mean distance (Å) | N positions | Mean distance (Å) |
| Substitution model with independent effect sizes (Eq 4). | Yes | 1 | 4.1 | 106 | 11.0 |
|  | No | 1 | 5.2 | 135 | 10.8 |
| Substitution model with effect sizes linked to position (Eq 5). | Yes | 6 | 0.9 | 113 | 12.3 |
|  | No | 4 | 0.6 | 110 | 15.3 |
| Structure aware model – position dependent effect sizes for substitutions and structure influencing inclusion probability (Eq 6). | Yes | 14 | 1.2 | 130 | 16.6 |
|  | No | 16 | 1.0 | 91 | 20.9 |

<sup>a</sup>Total number of variable positions evaluated = 199.

